# Mapping Mesoscale Axonal Projections in the Mouse Brain Using A 3D Convolutional Network

**DOI:** 10.1101/812644

**Authors:** Drew Friedmann, Albert Pun, Eliza L Adams, Jan H Lui, Justus M Kebschull, Sophie M Grutzner, Caitlin Castagnola, Marc Tessier-Lavigne, Liqun Luo

## Abstract

The projection targets of a neuronal population are a key feature of its anatomical characterization. Historically, tissue sectioning, confocal microscopy, and manual scoring of specific regions of interest have been used to generate coarse summaries of mesoscale projectomes. We present here TrailMap, a 3D convolutional network for extracting axonal projections from intact cleared mouse brains imaged by light-sheet microscopy. TrailMap allows region-based quantification of total axon content in large and complex 3D structures after registration to a standard reference atlas. The identification of axonal structures as thin as one voxel benefits from data augmentation but also requires a loss function that tolerates errors in annotation. A network trained with volumes of serotonergic axons in all major brain regions can be generalized to map and quantify axons from thalamocortical, deep cerebellar, and cortical projection neurons, validating transfer learning as a tool to adapt the model to novel categories of axonal morphology. Speed of training, ease of use, and accuracy improve over existing tools without a need for specialized computing hardware. Given the recent emphasis on genetically and functionally defining cell types in neural circuit analysis, TrailMap will facilitate automated extraction and quantification of axons from these specific cell types at the scale of the entire mouse brain, an essential component of deciphering their connectivity.

## Introduction

Volumetric imaging to visualize neurons in intact mouse brain tissue has become a widespread technique. Light-sheet microscopy has improved both the spatial and temporal resolution for live samples (Bouchard et al., 2015; Liu et al., 2018) while advances in tissue clearing have lowered the barrier to imaging intact organs and entire organisms at cellular resolution (Ariel, 2017; Pan et al., 2016). Correspondingly, there is a growing need for computational tools to analyze the resultant large datasets in three dimensions. Tissue clearing methods such as CLARITY and iDISCO have been successfully applied to the study of neuronal populations in the mouse brain and automated image analysis techniques have been developed for these volumetric datasets to localize and count simple objects, such as cell bodies of a given cell type or nuclei of recently active neurons (Chung et al., 2013; Menegas et al., 2015; Renier et al., 2016; Richardson and Lichtman, 2015). However, there has been less progress in software designed to segment and quantify axonal projections at the scale of the whole brains.

As single-cell sequencing techniques continue to dissect heterogeneity in neuronal populations (eg. Tasic et al., 2018; Zeisel et al., 2018) and as more genetic tools are generated to access these molecularly or functionally defined subpopulations, anatomical and circuit connectivity characterization is crucial to inform functional experiments (Luo et al., 2018; Sun et al., 2019). Traditionally and with some exceptions (eg. Oh et al., 2014; Zingg et al., 2014), ‘projectome’ analysis entails qualitatively scoring the density of axonal fibers in manually defined subregions selected from representative tissue sections and imaged by confocal microscopy. This introduces biases from the experimenter, including which thin tissue section best represents a large and complicated 3D brain region, how to bin axonal densities into high, medium, and low groups, whether to consider thin and thick axons equally, and whether to average or ignore density variation within a target region. Additionally, it can be difficult to precisely align these images to a reference atlas (Fürth et al., 2017). Volumetric imaging of stiff cleared samples and 3D registration to the Allen Brain Institute’s Common Coordinate Framework (CCF) has eliminated the need to select individual tissue sections (Allen Institute for Brain Science, 2017). However, without a computational method for quantifying axon content, researchers must still select and score representative optical sections (Schneeberger et al., 2019).

The automated identification and segmentation of axons from 3D images should circumvent these limitations. Recent application of deep convolutional neural networks (DCNNs) and Markov random fields to biomedical imaging have made excellent progress at segmenting grayscale CT and MRI volumes for medical applications (Alegro et al., 2017; Dong et al., 2018; Frasconi et al., 2014; Mathew et al., 2015; Thierbach et al., 2018). Other fluorescent imaging strategies including light-sheet, fMOST, and serial two-photon tomography have been combined with software like TeraVR, Vaa3D, Ilastik, and NeuroGPS-Tree to trace or otherwise reconstruct individual neurons (Peng et al., 2010; Quan et al., 2016; Wang et al., 2019; Winnubst et al., 2019; Zhou et al., 2018). However, accurate reconstruction often requires sparse labeling, but datasets with low cell numbers do not capture rare collateralization patterns.

As one of the most successful current DCNNs, the U-Net architecture has been used for local tracing of neurites in small volumes and for whole-brain reconstruction of brightly labeled vasculature (Çiçek et al., 2016; Falk et al., 2019; Di Giovanna et al., 2018; Gornet et al., 2019; Ronneberger et al., 2015). We posited that a 3D U-Net would be well suited for identifying axons, a similarly regular structural element albeit with a much lower signal-to-noise ratio, dramatic class imbalance, uneven background in different brain regions, and difficult annotation strategy. In addition to these challenges, whole-brain imaging of axons necessitates the inclusion of artifacts that contaminate the sample. The paired clearing and analysis pipeline we present here mitigates the impact of autofluorescent myelin and non-specific antibody labeling that interfere with the detection of thin axons. Our network, TrailMap (<u>T</u>issue <u>R</u>egistration and <u>A</u>utomated Identification of <u>L</u>ight-sheet <u>M</u>icroscope <u>A</u>cquired <u>P</u>rojectomes), provides a solution to all of these challenges. We demonstrate its generalization to multiple labeling strategies, cell types, and target brain regions. Alignment to the Allen Institute reference atlas allows for visualization and quantification of individual brain regions. We have also made available our best trained model such that any researcher with mesoscale projections in cleared brains can use TrailMap to process their image volumes or, with some additional training and transfer learning, adapt the model to their data.

## Results

### TrailMap procedures

To generate training data, we imaged 18 separate intact brains containing fluorescently labeled serotonergic axons (Fig. S1*A*). From these brains we cropped 36 substacks with linear dimensions of 100–300 voxels (Fig. 1*A*). Substacks were manually selected to represent the diversity of possible brain regions, background levels, and axon morphology. As with all neural networks, quality training data is crucial. Manual annotation of complete axonal structures in three dimensions is difficult, imprecise, and laborious for each volume to be annotated. One benefit of convolutional neural networks is the ability to input sparsely labeled 3D training data. We annotated 3–10 individual XY planes within each substack, at a spacing of 80–180 μm between labeled slices (Fig. 1*A*). Drawings of axons included single voxels representing the cross section of a thin axon passing through the labeled slice. In the same XY slice, a second label surrounding the axon annotation (“edges”) was automatically generated and the remaining unlabeled voxels in the slice were given a label for “background.” Unannotated slices remained without a label as previously indicated.

**Fig. 1.**
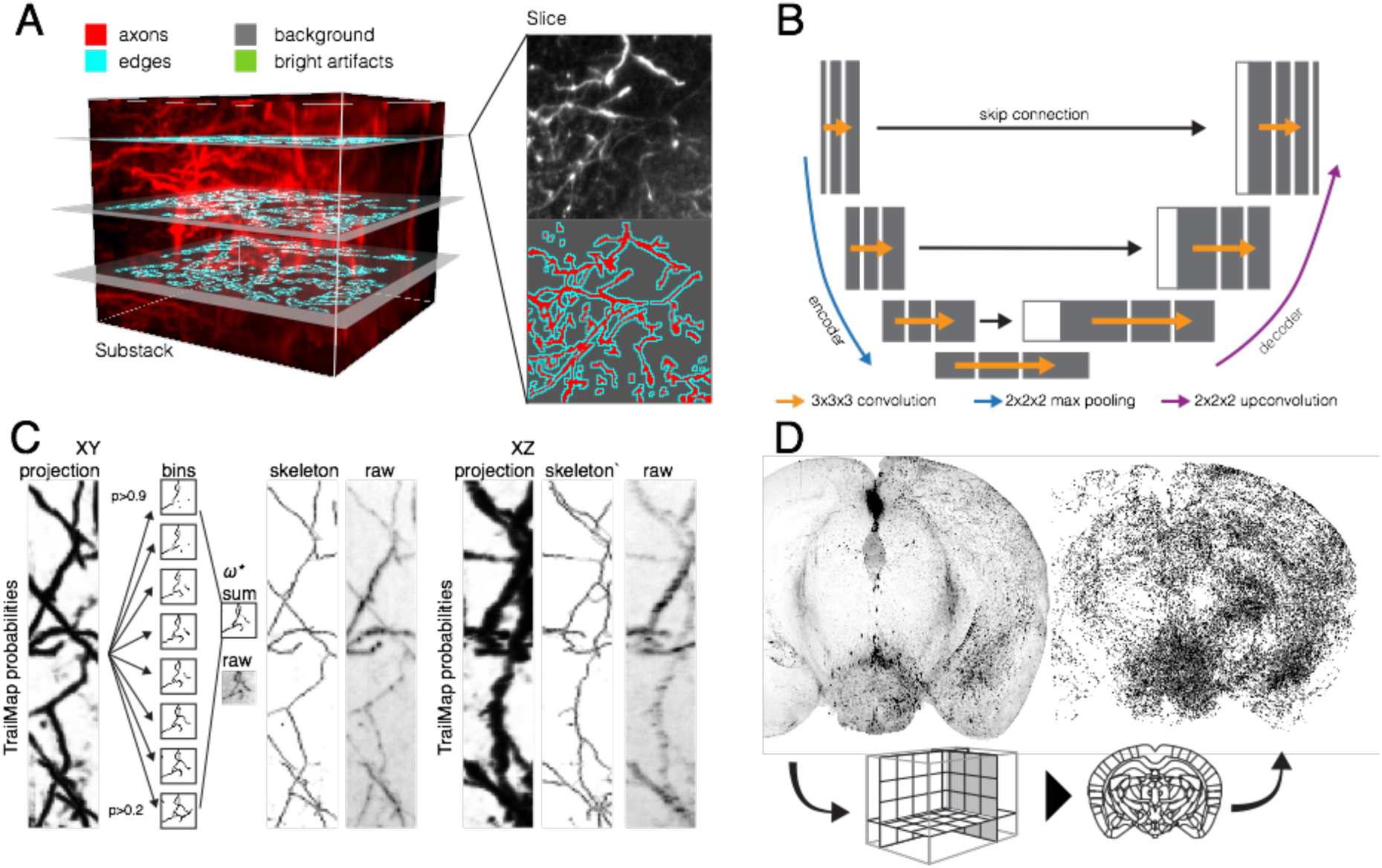
Overview of the TrailMap workflow to extract axonal projections from volumetric data. (*A*) Annotation strategy for a single subvolume (120×120×101 voxels). Three planes are labeled with separate hand-drawn annotations for background, artifacts, and axons. The 1-pixel width ‘edges’ label is automatically generated. (*B*) Basic architecture of the encoding and synthesis pathways of the 3D convolutional U-Net. Feature maps are shown in gray with the concatenated features in white. (*C*) A network output thinning strategy produces skeletons faithful to the raw data but with grayscale intensity reflecting probability rather than signal intensity or axon thickness. XY and XZ projections of one subvolume are shown (122×609×122 µm). (*D*) A 2 mm-thick volumetric coronal slab, before and after the TRAILMAP procedure, which includes axon extraction, skeletonization, and alignment to the Allen Brain Atlas Common Coordinate Framework.

Training data were derived from serotonin axons labeled by multiple strategies to better generalize the network (Fig. S1*A*). As serotonin neurons innervate nearly all forebrain structures, they provide excellent coverage for examples of axons in regions with variable cytoarchitecture and therefore, variable background texture and brightness. Our focus on imaging intact brains required the inclusion of contaminating artifacts from the staining and imaging protocol since these non-specific and bright artifacts are common in cleared brains and interfere with methods for axon identification. We addressed this in two ways, first by implementing a modified version of the AdipoClear tissue clearing protocol (Branch et al., 2019; Chi et al., 2018) that reduces the autofluorescence of myelin. As fiber tracts composed of unlabeled axons share characteristics with the labeled axons we aim to extract, this reduces the rate of false positives in structures such as the striatum (Fig. S1*B*). Second, we included 40 substacks containing representative examples of imaging artifacts and non-specific background and generated a fourth annotation label, “artifacts,” for these structures.

From this set of 76 substacks, we cropped and augmented 10,000 separate cubes of 64×64×64 voxels to use as the training set. Our validation set comprised 1,700 cubes extracted from 17 separate substacks, each cropped from one of nine brains not used to generate training data. The U-Net architecture included two 3D convolutions with batch normalization at each layer, 2×2×2 max pooling between layers on the encoder path, and 2×2×2 upconvolution between layers on the decoder path (Fig. 1*A*). Skip connections provide information necessary for recovering a high-resolution segmentation from the final 1×1×1 convolution. The final network was trained for 188 epochs over 20 hours, but typically reached a minimum in the validation loss approximately a third of the way into training. Subsequent divergence in the training and validation F1 scores (see Materials and Methods) indicated overfitting and as such, the final model weights were taken from the validation loss minimum (Fig. S1*C*–*E*).

For a given input cube, the network outputted a 36×36×36 volume containing voxel-wise axon predictions (0 < *p <* 1). Large volumes, including intact brains, were processed with a sliding window strategy. From this output, a thinning strategy was implemented to generate a skeletonized armature of the extracted axons (Fig. 1*C*). Grayscale values of the armature were the weighted sum of 3D skeletons generated from binarization of network outputs. For visualizations, this strategy maintained axon continuity across low probability stretches that would otherwise have been broken by a thresholding segmentation strategy. A separate benefit of this skeletonization strategy is that it treats all axons, thin and thick or dim and bright, equally for both visualization and quantification. A second imaging channel was used to collect autofluorescence, which in turn was aligned to the Allen Brain Atlas Common Coordinate Framework (CCF) via sequential linear and nonlinear transformation (http://elastix.isi.uu.nl/). These transformation vectors were then used to warp the axon armature into a standard reference space (Fig. 1*D*).

### Comparisons with random forest classifier

One of the most widely used tools for pixelwise classification and image segmentation is the random-forest-based software Ilastik (http://www.ilastik.org). We compared axon identification by TrailMap with multiple Ilastik classifiers and found TrailMap to be superior (best TrailMap model: recall: 0.752, precision: 0.377, 1-voxel exclusion zone precision: 0.790; best Ilastik classifier: recall: 0.208, precision: 0.661, 1-voxel exclusion zone precision: 0.867), but most notably in the clarity of the XZ projection (Fig. 2*A*-*C*). The increase in TrailMap’s precision when excluding ‘edge’ voxels suggested that many false positives are within one voxel of annotated axons. Ilastik examples used for comparison include classifiers trained in 3D with sparse annotations that include or exclude edge information as described for TrailMap, an example trained with images generated by iDISCO+, and a classifier trained in 2D using the exact set of labeled slices used to train TrailMap (Fig. S2*A*).

**Fig. 2.**
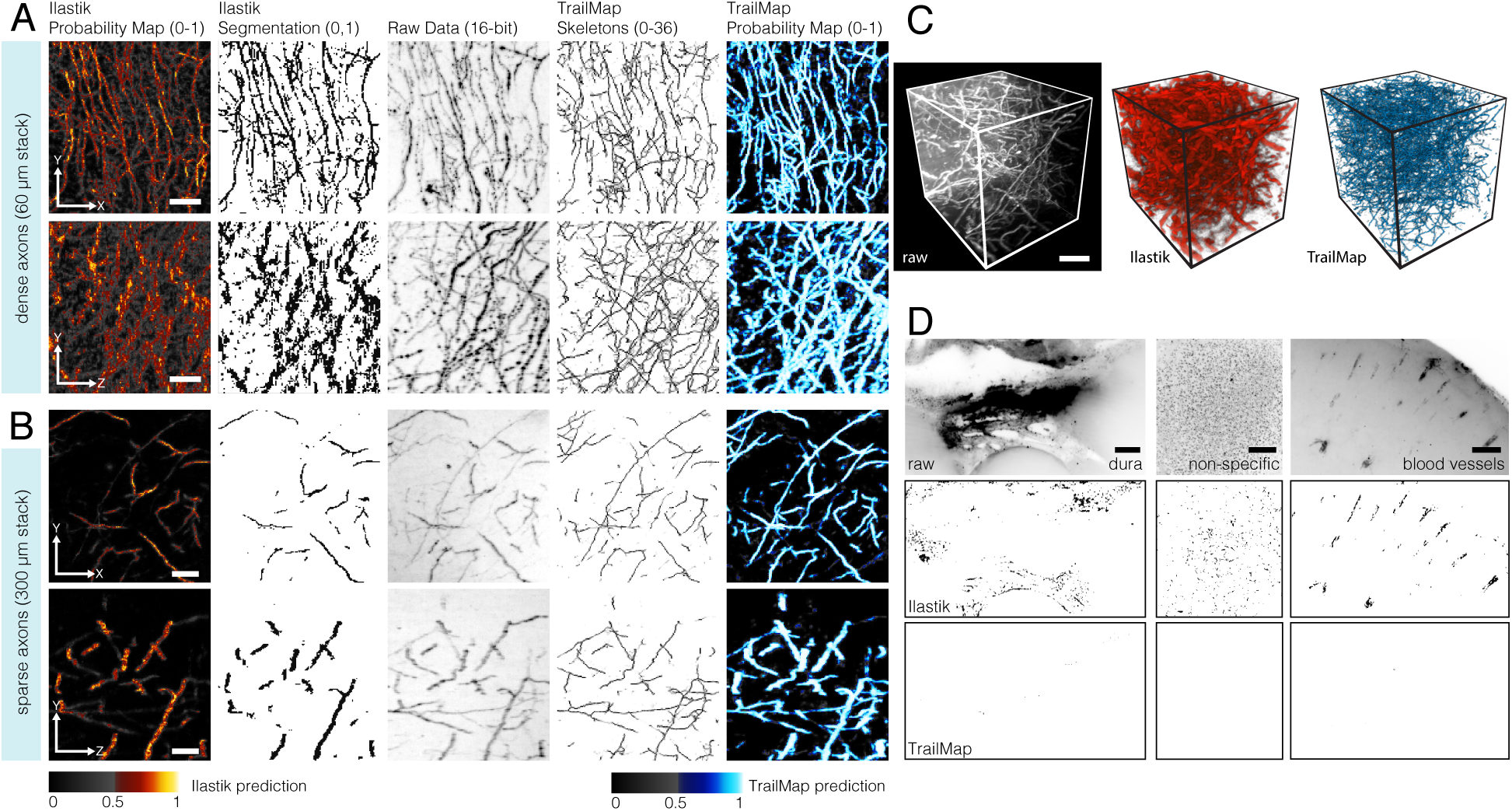
Comparison of TRAILMAP to a random forest classifier. (*A*) From left to right: 60 µm Z-projection of the probability map output of an Ilastik classifier trained on the same 2D slices as the TrailMap network; Typical segmentation strategy, at a p > 0.5 cutoff; Raw data for comparison; Skeletonized axonal armature extracted by TrailMap; Probability map output of the TrailMap network. To better indicate where *p* > 0.5, colormaps for images are grayscale below this threshold and colorized above it. Scale bar, 100 µm. Second row shows the same region as above, rotated to a X-projection. (*B*) Sparse axon identification by TrailMap and Ilastik, images show 300 µm of Z- or X-projection. Scale bar, 100 µm. (*C*) 3D maximum intensity projection of a raw volume of axons, the resultant Ilastik probabilities, and TrailMap skeletonization. Scale bar 40 μm. (*D*) Network output from examples of contaminating artifacts from non-specific antibody labeling, similar to those included in the training set. Raw data, Ilastik segmentation and TrailMap skeletonization are shown. Scale bar 200 μm.

To better understand which aspects of the TrailMap network contribute to its function, we trained multiple models without key components for comparison. As with all neural networks, we identified a number of hyperparameters that affected the quality of the network output. The most important was the inclusion of a weighted loss function to allow for imperfect annotations of thin structures. The cross-entropy calculation for voxels defined as “edges” were valued 30x less in calculating loss than the voxels annotated as axons. Manually labeling structures with a width of a single pixel introduced a large amount of human variability (Fig. S2*B*) and de-emphasizing the boundaries of these annotations allowed the network’s axon prediction to ‘jitter’ without producing unnaturally thick predictions (Fig. S2*C*). Relatedly, the total loss equation devalued background (7.5x) and artifact examples (1.875x) with respect to axon annotations—a set of weights that balances false positives and negatives and compensates for class imbalances in the training data (Fig. 2*D*, Fig. S2*C*, Table S1).

### Details of axonal projections

To determine TrailMap’s ability to identify axons from intact cleared samples, we tested a whole brain containing axon collaterals of serotonin neurons projecting to the regions of bed nucleus of stria terminalis. With the extracted axonal projectome transformed into the Allen Institute reference space, the axon armature could be overlaid on a template to better highlight their presence, absence, and structure in local subregions (Fig. 3*A*, Fig. S3). However, it was difficult to resolve steep changes in density or local hotspots of innervation without selectively viewing very thin sections. By averaging axons with a rolling sphere filter, a small hyper-dense zone is revealed in amygdala that would have been missed in region-based quantifications (Fig. 3*B*, arrow). Total axon content in large complicated brain regions could be quantified (Ren et al., 2019) or otherwise projected and visualized to retain local density information in three dimensions (Fig. 3*C* and *D*).

**Fig. 3.**
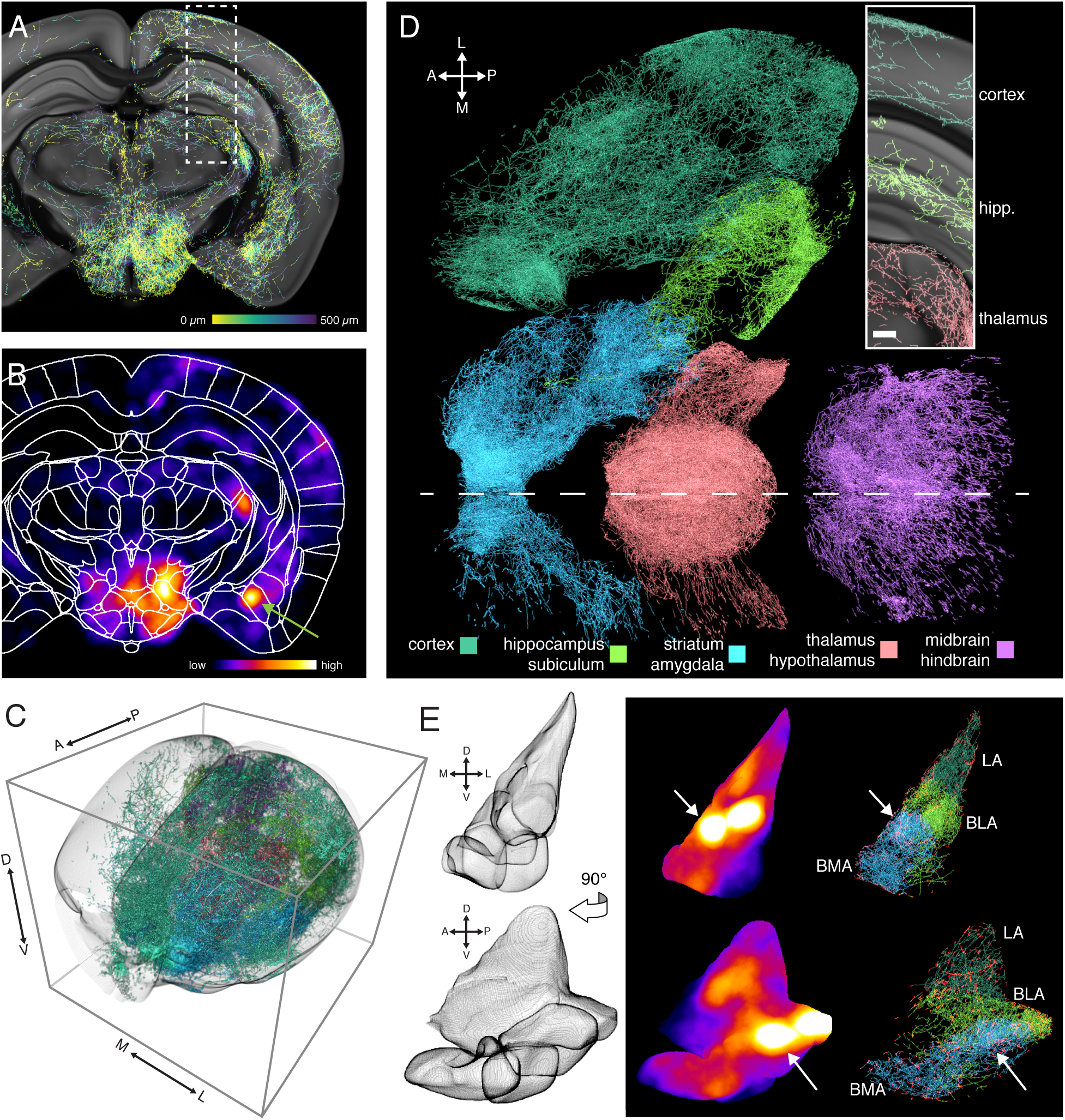
Volumetric visualizations highlight patterns of axonal innervation. (*A*) Coronal Z-projection of extracted serotonergic axons, color coded by depth (0–500 µm) and overlaid on the CCF-aligned serial two-photon reference atlas. See Fig. S1 for viral-transgenic labeling strategy. (*B*) The same volumetric slab as in *A* presented as a density heatmap calculated by averaging a rolling sphere (radius = 225 μm). Green arrow highlights a density hotspot in the amygdala. (*C*) TrailMap-extracted serotonergic axons innervating forebrain are subdivided and color coded based on their presence in ABA-defined target regions. (*D*) Same brain as in *C* as seen from a dorsal viewpoint, with major subdivisions spatially separated. Midline represented by a dashed white line. Inset highlights the region indicated by the dashed box in *A*, Z-projection 500 μm, scale bar 200 μm. (*E*) Left, mesh shell armature of the combined structures of the lateral (LA), basolateral (BLA), and basomedial (BMA) amygdala in coronal and sagittal views. Right, density heatmap and extracted axons of the amygdala for the same views. LA, BLA, and BMA are color coded in shades of green/blue by structure and axonal entry/exit points to the amygdala are colored in red. White arrows highlight the same density hotspot indicated in *B*.

An added benefit of the transformation into a reference coordinate system is that brain regions as defined by the Allen Institute could be used as masks for highlighting axons in individual regions of interest. As an example, the aforementioned density observed in amygdala is revealed to be contributed by two nearby dense zones in the basomedial and basolateral amygdala, respectively (Fig. 3*E*).

### Generalization to other types of neurons and brain regions

Given that TrailMap was trained exclusively on serotonergic neurons, it may not generalize to other cell types if their axons are of different sizes, tortuosity, or bouton density. The network may also fail if they are labeled with a different genetic strategy or imaged at different magnification. However, our data augmentation provided enough variation in training data to produce excellent results in extracting axons from multiple additional cell types, fluorophores, viruses, and imaging setups. TrailMap successfully extracted brain-wide axons from pons-projecting deep cerebellar nuclei neurons retrogradely infected with AAV_retro_-Cre and locally injected in each DCN with AAV-DIO-tdTomato (Fig. 4*A*). Visualizing the whole brain (Fig. 4*A*) or the thalamus (Fig. 4*B*) in 3D reveals the contours of the contralateral projections of these neurons without obscuring information at the injection site or ipsilateral targets.

**Fig. 4.**
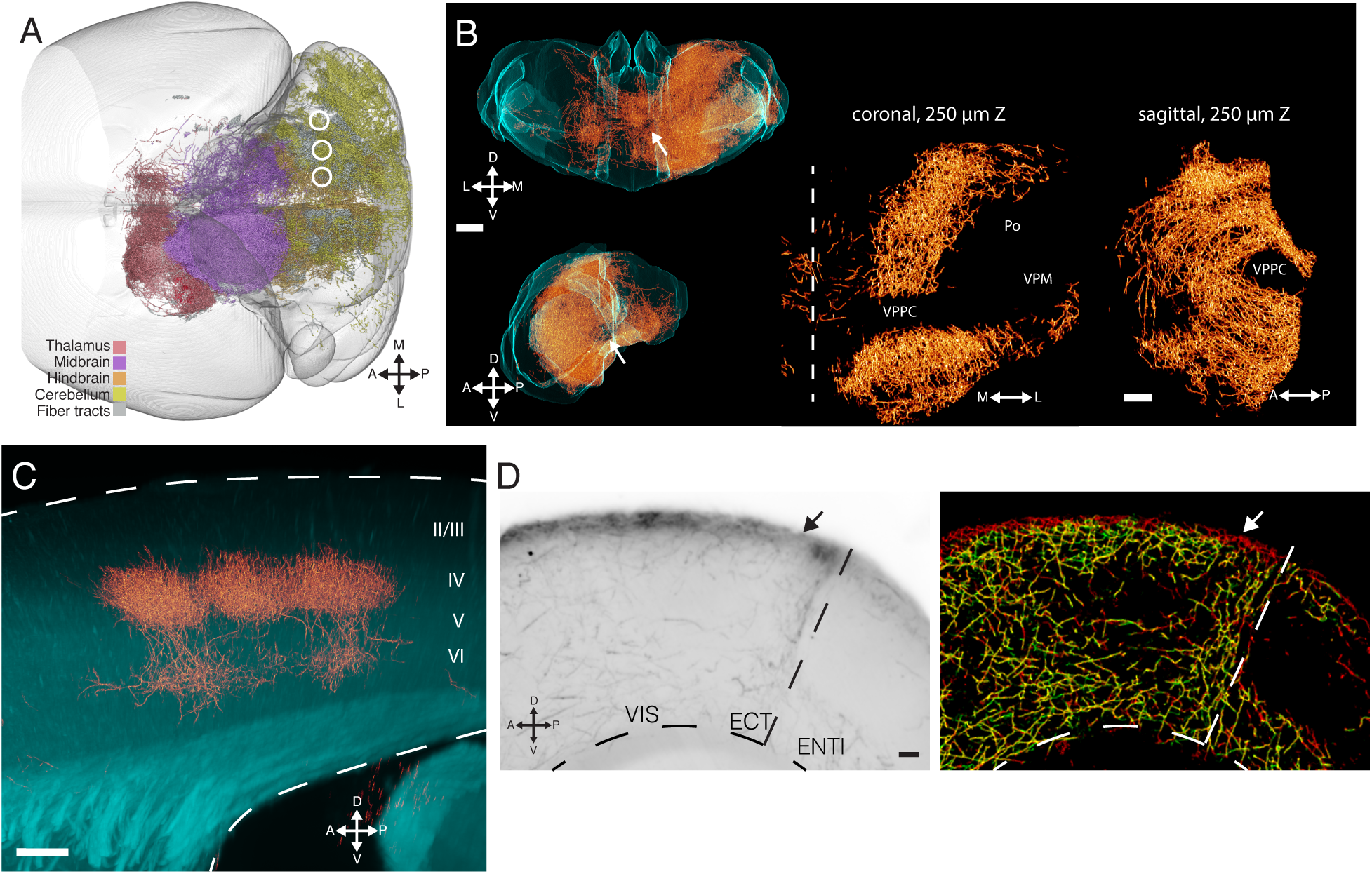
Generalization to other cell types with and without transfer learning. (*A*) Dorsal view of posterior brain with extracted axon collaterals from pons-projecting deep cerebellar nuclei neurons color coded by their presence in major subregions. Injection sites in the right hemisphere lateral, interposed, and medial DCN from the same brain are indicated. (*B*) Coronal (top) and sagittal (bottom) views of the extracted axons (orange) within the structure of thalamus (cyan) from the brain in *A*. Scale bar 500 μm. Zoomed images (right) of midline thalamus show exclusion of axons from specific subregions. VPPC: ventral posterior nucleus, parvicellular part; VPM:ventral posteriomedial nucleus; and Po: posterior thalamic nuclear group. Midline shown as dashed vertical line; scale bar, 200 μm. (*C*) Extracted thalamocortical axons in barrel cortex. XZ-projection of a 3D volume extracted from a flatmount imaged cortex, rotated and aligned to barrel rows. Dashed lines indicate upper and lower bounds of cortex. Z projection 60 μm; scale bar, 100 µm. (*D*) Left, raw image of axons of prefrontal cortex neurons in posterior cortex. VIS, visual cortex; ECT, ectorhinal; ENTl, lateral entorhinal. Right, axons extracted by TrailMap model before (green) and after (red) transfer learning. Z-projection 80 μm; scale bar, 100 μm. Labeling procedure, virus, and transgenic mouse for all panels are described in Fig. S1 and Methods.

We also tested TrailMap on functionally identified thalamocortical axons in barrel cortex labeled by conditional AAV-DIO-CHR2-mCherry and TRAP2 activity-dependent expression of Cre recombinase (DeNardo et al., 2019). The localized expression and higher density of these axons necessitates imaging at higher magnification, resulting in a change in axon appearance in the imaging volume. Buoyed by the spatial scaling included in the data augmentation, TrailMap reliably revealed the dense thalamic innervation of somatosensory layer IV and weaker innervation of layer VI (Fig. 4*C*). Notably, though Ilastik performs moderately well on serotonergic axons, it fails to predict thalamocortical axons, perhaps owing to the change in scale of the imaging strategy (Fig. S4).

Finally, TrailMap also extracted cortico-cortical projection axons from prefrontal cortex (PFC) labeled by retrograde CAV-Cre and AAV-DIO-mGFP-2A-synaptophysin-mRuby. Axons from PFC neurons imaged in posterior visual and entorhinal cortices were identified with the exception of the most superficial axons in layer I (Fig. 4*D*). The failure to identify layer I axons could be because the serotonergic training set did not include examples of superficial axons; as a result, the trained network used the presence of low intensity grayscale values outside the brain to influence the prediction for each test cube containing the edge of the sample. Using 17 new training substacks from brains with annotated superficial axons from PFC cortical projections and 5 new validation volumes, we performed transfer learning using our best model as the initial weights. After just five epochs, the model successfully identified these layer I axons (Fig. 4*D*).

## Discussion

Here we present an adaptation of a 3D U-Net tuned for identifying axonal structures within noisy whole-brain volumetric data. Our trained network, TrailMap, is specifically designed to extract mesoscale projectomes rather than reconstructions of individual neurons. For intact brains with hundreds of labeled neurons or zones of high-density axon terminals, we are not aware of a computational alternative that can reliably identify these axons. Our clearing and image processing pipeline address a number of challenges that have prevented these mesoscale analyses up till now. First, all clearing techniques have at least some issues with background or non-specific labeling that can interfere with automated image analysis. Removing myelinated fiber tracts with a modified AdipoClear protocol greatly improved TrailMap’s precision in structures such as the striatum. Second, our weighted loss function considers manually annotated, non-specific, bright signals separately from other background areas—an essential step in reducing false positives. Relatedly, by devaluing the loss calculated for voxels adjacent to axon annotations, we reduced the rate of false negatives by allowing the network to err by deviations of a single voxel. Third, we present a strategy for thinning TrailMap’s output to construct an armature of predicted axons. This armature improves visualizations and reduces biases in analysis and quantification by giving each axon equal thickness independent of staining intensity or imaging parameters.

Aligning armatures and density maps to the Allen Institute’s reference brain highlights which brain regions are preferentially innervated or avoided by a specific projection (Ren et al., 2019). Given that some brain regions are defined with less certainty, it will be possible to use the axon targeting of specific cell types to refine regional boundaries. As TrailMap can separate thalamocortical projections to individual whisker barrels (Fig. 4*C*), it will be interesting to locate other areas with sharp axon density gradients that demarcate substructures within larger brain regions. These collateralization maps will also assist neuroanatomists investigating the efferent projection patterns of defined cell populations. TrailMap has the added benefit of 3D, intact structures as the basis for quantification, but also the ability to process samples and images in parallel, reducing the active labor required to generate a complete dataset that spans the entire brain.

TrailMap code is publicly available, along with the weights for our best model and example data. A consumer-grade GPU is sufficient for both processing samples and for performing transfer learning, while training a new model from scratch benefits from the speed and memory availability from cloud computing services. We hope that TrailMap’s ease of use will lead to its implementation by users as they test brain clearing as a tool to visualize their neurons of interest. We expect that as neuronal cell types are becoming increasingly defined by molecular markers (Jorgenson et al., 2015), TrailMap can be used to map and quantify whole-brain projections of these neuronal types using viral-genetic and intersectional genetic strategies (eg. Ren et al., 2019).

## Materials and Methods

### Animals

All animal procedures followed animal care guidelines approved by Stanford University’s Administrative Panel on Laboratory Animal Care (APLAC). Individual genetic lines include wildtype mice of the C57BL/6J and CD1 strains, TRAP2 (*Fos-iCreER*^*T2*^) available from Jackson, Stock # 030323), Ai65 (Jackson, Stock # 021875), and *Sert-Cre* (MMRRC, Stock #017260-UCD). Mice were group-housed in plastic cages with disposable bedding on a 12 hours light/dark cycle with food and water available ad libitum.

### Viruses

Combinations of transgenic animals and viral constructs are outlined in Fig. S1. Viruses used to label serotonergic axons include: AAV-DJ-hSyn-DIO-HM3D(Gq)-mCherry (Stanford vector core), AAV8-ef1α-DIO^FRT^-loxp-STOP-loxp-mGFP (Stanford vector core; Ren et al. *Cell* 2018), AAV-retro-CAG-DIO-Flp (Salk Institute GT3 core; Ren et al. *Cell* 2018), AAV8-CAG-DIO-tdTomato (UNC vector core, Boyden group), AAV_retro_-ef1α-Cre (Salk Institute GT3 core), AAV8-ef1α-DIO-CHR2-mCherry (Stanford vector core), AAV8-hSyn-DIO-mGFP-2A-synaptophysin-mRuby (Stanford vector core, Addgene #71760), and Cav-Cre (Eric Kremer; (Schwarz et al., 2015)).

### AdipoClear labeling and clearing pipeline

Mice were transcardially perfused with 20 ml 1x PBS containing 10 μg/μl heparin followed by 20 ml ice cold 4% PFA and post fixed overnight at 4° C. All steps in the labeling and clearing protocol are on a rocker at room temperature for a 1-hour duration unless otherwise noted. Brains are washed 3x in 1x PBS and once in B1n before dehydrating stepwise into 100% methanol (20, 40, 60, 80% steps). Two additional washes in 100% methanol remove all the water before an overnight incubation in 2:1 dichloromethane (DCM):methanol. The following day, 2 washes in 100% DCM and 3 washes in methanol precede 4 hours in a 5:1 methanol:30% hydrogen peroxide mixture. Stepwise brains are rehydrated into B1n (60, 40, 20% methanol), washed once in B1n and then permeabilized 2x in PTxwH containing 0.3M glycine and 5% DMSO. Samples are washed 3x in PTxwH before adding primary antibody (chicken anti-GFP, 1:2000, Aves Labs; rabbit anti-RFP, 1:1000, Rockland Inc). Incubation is for 7-11 days rocking at 37° C, subsequent washes also at this temperature. Brains are washed 5x in PTxwH over 12 hours and then 1x each day for 2 additional days. Secondary antibody (donkey anti-rabbit, 1:1000, Thermo; donkey anti-chicken, 1:2000, Jackson) is incubated rocking at 37° C for 5-9 days. Wash 5x in PTxwH over 12 hours and then 1x each day for 2 additional days. Samples are dehydrated stepwise into methanol, as before, but with water as the counterpart, then washed 3x in 100% methanol, overnight in 2:1 DCM:methanol, and 2x 100% DCM the next morning. Extend the second wash in DCM until the brain sinks. Transfer to dibenzyl ether (DBE) in a fresh tube and incubate rocking for 4 hours before storing in another fresh tube of DBE at room temperature. *Solutions--*B1n: 1:1,000 Triton X-100, 2% w/v glycine, 1:10,000 NaOH 10N, 0.02% sodium azide. PTxWH: in 1x PBS, 1:1,000 Triton X-100, 1:2,000 Tween-20, 2 μg/μl heparin, 0.02% sodium azide.

### Light-sheet imaging

Image stacks were acquired with the LaVision Ultramicroscope II light-sheet microscope using the 2x objective at 0.8x optical zoom (4.0625 μm/pixel, XY dimension). Thalamocortical axons imaged at 1.299 μm/voxel, XY dimension. Maximum sheet objective NA combined with 20 steps of horizontal translation of the light-sheet improves axial resolution. Z-step size 3 μm. Axon images were acquired with the 640 nm laser and a partner volume of autofluorescence of equal dimensions was acquired with the 488 nm laser. No pre-processing steps were taken before entering the TrailMap pipeline.

### TrailMap annotation strategy

To create the training set for axons, we used sparse annotations to label volumes of approximately 100–300 voxels/side. These were cropped from 18 samples across experimental batches from each of the three serotonergic neuron labeling strategies outlined in Fig. S1. Two experts traced axons in these 36 different substacks by labeling single XY-planes from the stack every ∼20–30 slices. Additionally, 40 examples of bright artifacts were found from these volumes and labeled as artifacts through an intensity thresholding method. Some examples contained both manually annotated axons and thresholded labels for artifacts. From these labeled volumes, 10,000 training examples were generated by cropping cubes (linear dimensions, 64×64×64) from random locations within each labeled substack. We also introduced an additional independent label, referenced as “edges.” This label was programmatically added to surround the manually drawn “axon” label in the single-voxel thick XY plane. This label was generated and subsequently given less weight in the loss function specifically to help the network converge by reducing the penalty for making off-by-one errors in voxels next to axons. Both edge and artifact labels are only used for weighting in the loss function, but are still counted as background. We created a validation set using the same methodology as the training set; however, to test for resilience against the potential impacts of staining, imaging, and other technical variation, the substacks used for the validation set were from different experimental batches than the training set.

### Data augmentation and network structure

Due to the simple cylindrical shape of an average axon segment, Z-score normalization was avoided as it removed the raw intensity information from the original image volume. Without this information, the network could not differentiate natural variability in the background from true axons. To provide robustness to signal intensity variation during training, chunks were augmented in real time through random scaling and summating random constants. We used a 3D-U-Net architecture (Çiçek et al., 2016) with input size 64^3^ and output of a 36^3^ segmentation. The output dimensions are smaller, accounting for the lack of adequate information at the perimeter of the input cube to make an accurate prediction. To segment large volumes of intact brains, the network was applied in a non-overlapping sliding window fashion.

We used a binary cross entropy pixel-wise loss function, where axons, background, edges, and artifacts were given static weights to compensate for class imbalances and the structural nature of the axon. The binary loss function is calculated as (−(*y*log(*p*)+(1−*y*)log(1−*p*)))\**w* for every voxel with the predicted value (*p*), the true label (*y*), and a weight (*w*) determined from the true label’s associated weight. The network was trained on Amazon Web Services using p3.xlarge instances for approximately 16 hours. Training is approximately 2x slower, using 6 volumes/batch (as compared to 8 volumes/batch on AWS) with a NVIDIA GeForce 11GB 1080 Ti GPU. We trained many models with varying weights for the loss function and data scaling factor and picked the model that resulted in the lowest validation loss.

### Model Evaluation

To evaluate the network, we compared the validation set output by the model to ground truth human annotations. Each pixel was classified to be TP (true positive), TN (true negative), FP (false positive), or FN (false negative). It is important to note that we did not include pixels which were labeled as edges because these were predetermined to be ambiguous cases and would not be an accurate representation of the network’s performance. Using these four classes, we used the following formulas for metrics; Precision = TP/(TP + FP); Recall = TP/(TP + FN); F1 Value = 2 * TP/(2*TP + FP + FN); Jaccard Index: TP / (FP + TP + FN). Human-human comparison was done by having two separate experts annotate the same 41 slices from each of 8 separate substacks before proceeding with the same set of evaluations.

### Skeletonization and 3D alignment to CCF

Probabilistic volumes from the output of TrailMap were binarized at eight separate thresholds, from 0.2 to 0.9. These binned logical volumes were skeletonized in three dimensions before being summed back together, weighted by their initial probability bin. The resulting armature thus retains information about TrailMap’s prediction confidence without breaking connected structures by threshold segmentation. Small, truncated, and disconnected objects were removed as previously described (Ren et al., 2018). We downsampled this armature from 4.0625 μm/pixel (XY) and 3 μm/pixel (Z) into a 5×5×5 μm space and also downsampled the autofluorescence image into a 25×25×25 μm space. The autofluorescence channel was aligned to the Allen Institute’s 25 μm reference brain acquired by serial two photon tomography. These affine and bspline transformation parameters were used to warp the axons into the CCF at 5 μm scaling. Once in the CCF, ABA region masks can be implemented for pseudocoloring, cropping, and quantifying axon content on a region-by-region basis. Rolling sphere voxelization with a radius of 45 voxels (225 μm) operates by summing total axon content after binarization.

## Acknowledgements

We thank Ethan Richman for helpful discussions and comments on the manuscript. D.F was supported by NINDS T32 NS007280, J.H.L. by NIMH K01 MH114022, J.M.K. by a Jane Coffin Childs postdoctoral fellowship, E.L.A. by the Stanford Bio-X Bowes PhD Fellowship. This work was supported by NIH Grant R01 NS104698 (to L.L.). L.L. is a Howard Hughes Medical Institute investigator.

