## Supplemental Figures for "Mapping Mesoscale Axonal Projections in the Mouse Brain Using A 3D Convolutional Network"

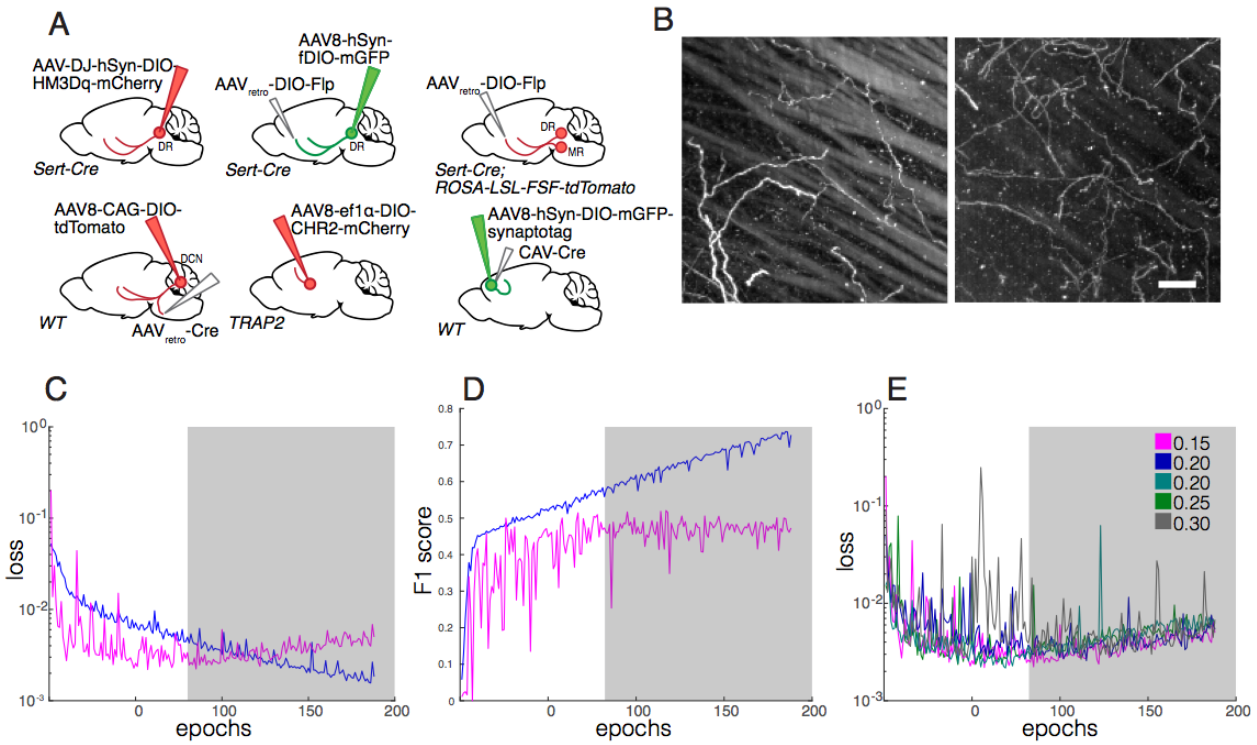


**Fig. S1.** Viral-genetic strategies for axon labeling, clearing methods, and specs for network training. (*A*) Viral-genetic strategies used for labeling axons used in training and testing. Top row: three serotonergic neuron labeling strategies used for creating the network training data. Top right, strategy used to generate the whole-brain data in Figure 3. Bottom row: additional strategies used to label axons for transfer learning training data and as seen in Figure 4 *A-B, C,* and *D* respectively. (*B*) Left, iDISCO+ clearing protocol does not remove autofluorescent fiber tracts in striatum while AdipoClear (right) performs better. Z-projection 285 μm; scale bar, 100 μm. (*C*) Training set (blue) and validation set (magenta) loss across 188 epochs. Gray shading indicates a positive slope in the validation loss and overfitting to the training set. (*D*) Same as in *C*, except showing the F1 score; see Materials and Methods for equations for metrics. (*E*) Validation loss for each of five training runs with variable intensity scaling factors during data augmentation. Without Z-score normalization of training data, we used a random scaling factor between zero and the depicted values. Higher values tended to increase the validation loss minimum. The magenta trace is the same as in *C* and *D*.

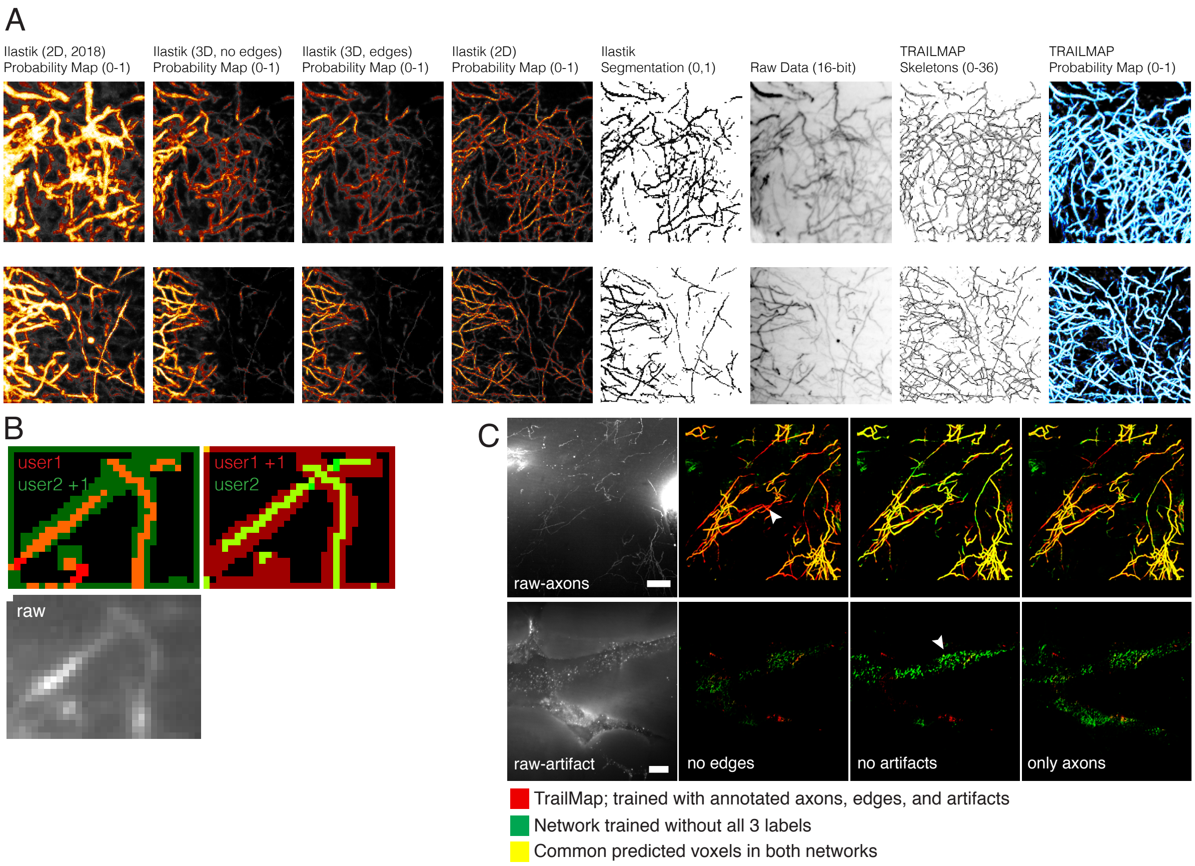


**Fig. S2.** Comparisons of TrailMap to Ilastik classifiers and models trained without all annotation categories. (*A*) Additional examples comparing four separate Ilastik classifiers with TrailMap. From left to right: 2D classifier, trained on iDISCO+ cleared brains (Ren et al., 2018); 3D classifier trained with only axon and background annotations; 3D classifier trained with axons, background, artifact, and edges annotations; 2D classifier trained with every slice of raw data used in the TrailMap training set; segmentation of the 2D Ilastik classifier at $p > 0.5$; raw data; TrailMap armatures; and TrailMap probabilities. Colormaps as in Main Figure 2. (*B*) Example annotations from two separate users. The bright pixels are the manual annotation and dim pixels are automatically generated for the edge label. User 2 labeled more conservatively, with 94.53% of their annotation falling within the User 1 + edges zone. 74.75% of User 1 annotations fall within the User 2 + edges zone. n = 8 substacks, 41 slices. (*C*) TrailMap output examples when excluding artifacts, edges, or both from the weighted loss equation. Arrowheads mark false negatives when excluding edges from the training data (top left) and false positives when excluding artifacts from training data (bottom center).

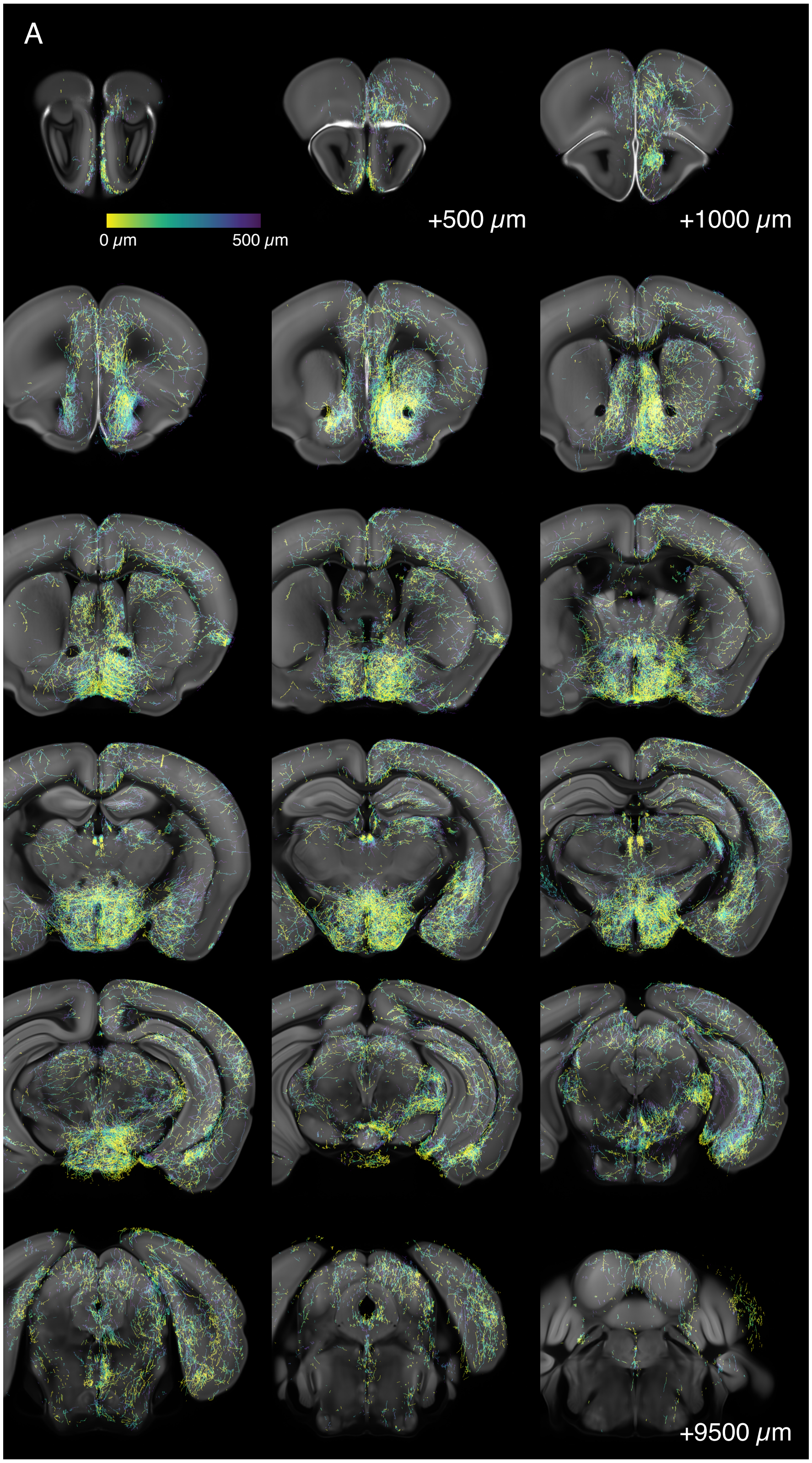

**Fig. S3**. Serotonergic axons across a whole brain as seen in Fig. 3A. Each panel represents 500 μm of Z-depth in the coronal axis, color-coded by depth as indicated in the top left.

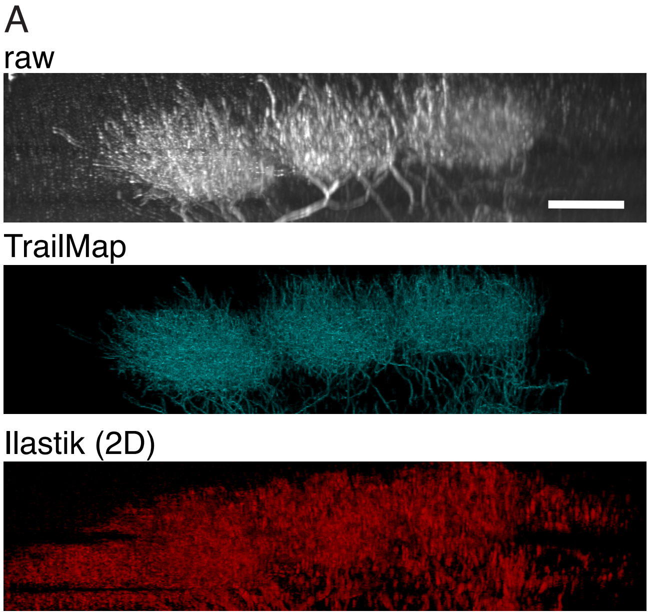


**Fig. S4** Thalamocortical axons as seen in Fig. 4C. Top, raw data; middle, TrailMap extracted armature; bottom, Ilastik (2D classifier) probability map. All images are XZ-projections, scale bar 100 μm.
